# Dual role of GABA_B_ receptor in oligodendrocyte function and immune modulation in experimental multiple sclerosis

**DOI:** 10.1101/2025.09.09.675094

**Authors:** Laura Bayón-Cordero, Blanca Isabel Ochoa-Bueno, Irene Luengas-Escuza, Rodrigo Senovilla, Izaskun Buendía, Chao Zheng, Eneritz Agirre, Gonçalo Castelo-Branco, Laura Amo, Francisco Borrego, Fernando García-Moreno, Frank Kirchhoff, Xianshu Bai, Carlos Matute, María Victoria Sánchez-Gómez

## Abstract

GABA_B_ receptors (GABA_B_R) mediate the actions of the inhibitory neurotransmitter GABA in the central nervous system, regulating key processes such as synaptic activity, interneuron communication and excitation-inhibition balance in the brain. Recent studies using the GABA_B_R agonist baclofen have revealed a critical role for these receptors in promoting oligodendroglial differentiation in both health and disease, highlighting their potential as therapeutic targets in demyelinating diseases such as multiple sclerosis (MS). In this study, we identify a dual role for oligodendroglial GABA_B_R in experimental autoimmune encephalomyelitis (EAE), an animal model of MS. Conditional deletion of the GABA_B1_ subunit in NG2+ cells ameliorates acute disease symptoms while inducing an immune-like phenotype in oligodendrocytes. This immunomodulatory role is further supported by pharmacological activation of GABA_B_R in oligodendrocytes, which reduces the expression of MHC class II in these cells. Notably, baclofen treatment after EAE symptom onset attenuates the course of the disease while enhancing oligodendrocyte progenitor cell differentiation and suppressing T cell infiltration into demyelinating lesions. Moreover, prophylactic baclofen administration delays disease onset and further decreases immune cell recruitment into the spinal cord, underscoring its potent immunomodulatory effect. These data demonstrate that GABA_B_R signaling exerts context-dependent effects on both oligodendrocyte lineage progression and neuroinflammatory responses. Importantly, these compelling findings validate baclofen, a drug already approved for MS-associated spasticity, as a promising candidate for therapies targeting both inflammation and remyelination, advancing our understanding of glial-immune interactions in demyelinating diseases and supporting the translational potential of GABA_B_R modulation in MS.

## Introduction

Multiple sclerosis (MS) stands as one of the main causes of non-traumatic neurological disability in young adults. It is a chronic inflammatory, neurodegenerative and demyelinating disease of the central nervous system (CNS), affecting more than 2 million people worldwide (Yamout & Alroughani, 2018). Immunopathogenesis in MS is driven by immune attack against CNS antigens, primarily through the action of myelin-reactive T cells from the adaptive immune system, which ultimately cause myelin breakdown, oligodendrocyte injury and axonal degeneration (Dendrou et al., 2015; Garg & Smith, 2015; Molina-Gonzalez et al., 2022). Thus, current strategies against MS target the inflammatory component of the disease, modulating features such as immune cell activity, cytokine secretion or lymphocyte infiltration into the CNS. These are highly effective in preventing the appearance of relapsing episodes, but their efficacy diminishes in progressive stages of the disease, raising awareness about the need for new therapies. In this context, remyelination is a crucial reparative mechanism in MS, presenting high potential for restoring neurological function, repairing existing damage and enhancing neuroprotection (Franklin & ffrench-Constant, 2017). Consequently, strategies promoting remyelination emerge as promising alternatives that should be combined with immunomodulators for the treatment of MS, in order to tackle the two main aspects of MS: immune cell activation and demyelination.

In the CNS, oligodendrocyte progenitor cells (OPCs) differentiate into mature oligodendrocytes (MOLs) to ensure myelin regeneration in the context of MS (Simons & Nave, 2016). Our latest studies demonstrate that GABA_B_ receptor (GABA_B_R) activation enhances OPC differentiation *in vitro* (Serrano-Regal et al., 2020a), and that GABA_B_R selective agonist baclofen accelerates remyelination after lysolecithin-induced primary demyelination in the mouse spinal cord (Serrano-Regal et al., 2022). However, these approaches fail to establish the contribution of oligodendroglial GABA_B_R in the context of MS. Furthermore, lysolecithin models fail to recapitulate the complexity of MS and therefore are not effective in showing the real potential of baclofen as a strategy against progressive stages of the disease.

Here, we first investigate the contribution of oligodendroglial GABA_B_R during the course of the experimental autoimmune encephalomyelitis (EAE), an animal model of MS that mirrors both the acute and chronic phases of the disease. We uncover a novel function of GABA_B_Rs in shaping the immune phenotype in oligodendrocytes. Importantly, we identify a novel role of GABA_B_R in oligodendrocytes, modifying their immune phenotype. Moreover, we then explore the potential of GABA_B_R agonist baclofen, a drug already used in clinic to treat spasticity in patients with MS, as a protective treatment in the EAE model. Our findings underscore the significance of GABA_B_R in the modulation of immune responses and OPC differentiation in demyelinating diseases and identify its agonist baclofen as a promising candidate for therapeutic strategies in MS.

## Results

### GABA_B1_ deletion in OPCs reduces acute EAE severity

We previously identified GABA_B_R as a promising therapeutic target for demyelinating diseases (Serrano-Regal et al., 2020a; 2022). To further explore the therapeutic potential of GABA_B_R, we investigate the role of oligodendroglial GABA_B_R in OPC differentiation. For that, we examine the effects of GABA_B_R signaling disruption through gene silencing of GABA_B1_ subunit in cortical OPCs, thereby reducing receptor activity in these cells. Immunoblot analyses revealed moderate silencing efficiency at 3 days *in vitro* (DIV) following transfection with *Gabbr1* esiRNA (Fig. 1a). Additionally, this reduction in GABA_B1_ subunit expression was accompanied by a significant decrease in the expression of MBP (Fig. 1a), suggesting that even a small reduction in GABA_B1_ levels was sufficient to impair MBP expression. Next, we determined the relevance GABA_B_R in the EAE model, and so we induced EAE in WT and cKO NG2-CreERT2: GABA_B1_ ^fl/fl^;tdTomato mice (Fang et al., 2022). Recombination in the cKO theoretically leading to GABA_B1_ deletion was confirmed by tdTomato expression. Given the reduced expression of MBP following GABA_B1_ silencing *in vitro*, as well as the impairment in oligodendroglial differentiation observed after NG2-GABA_B1_ ablation during development (Fang et al., 2022), we initially expected that GABA_B1_ deletion in OPCs would be detrimental to the course of the disease. However, cKO mice exhibited significantly milder disease-associated symptoms during the acute phase of the model compared to WT mice, although this difference was later reversed during the chronic phase (Fig. 1b). Histological analysis of the spinal cord of WT and cKO mice at 20 dpi (peak of the symptoms) revealed a remarkable reduction in the percentage of white matter area occupied by the lesions (Fig. 1c). These results indicate that GABA_B1_ genetic deletion in NG2^+^ cells attenuates the course of the acute phase in the EAE.

**Figure 1.**
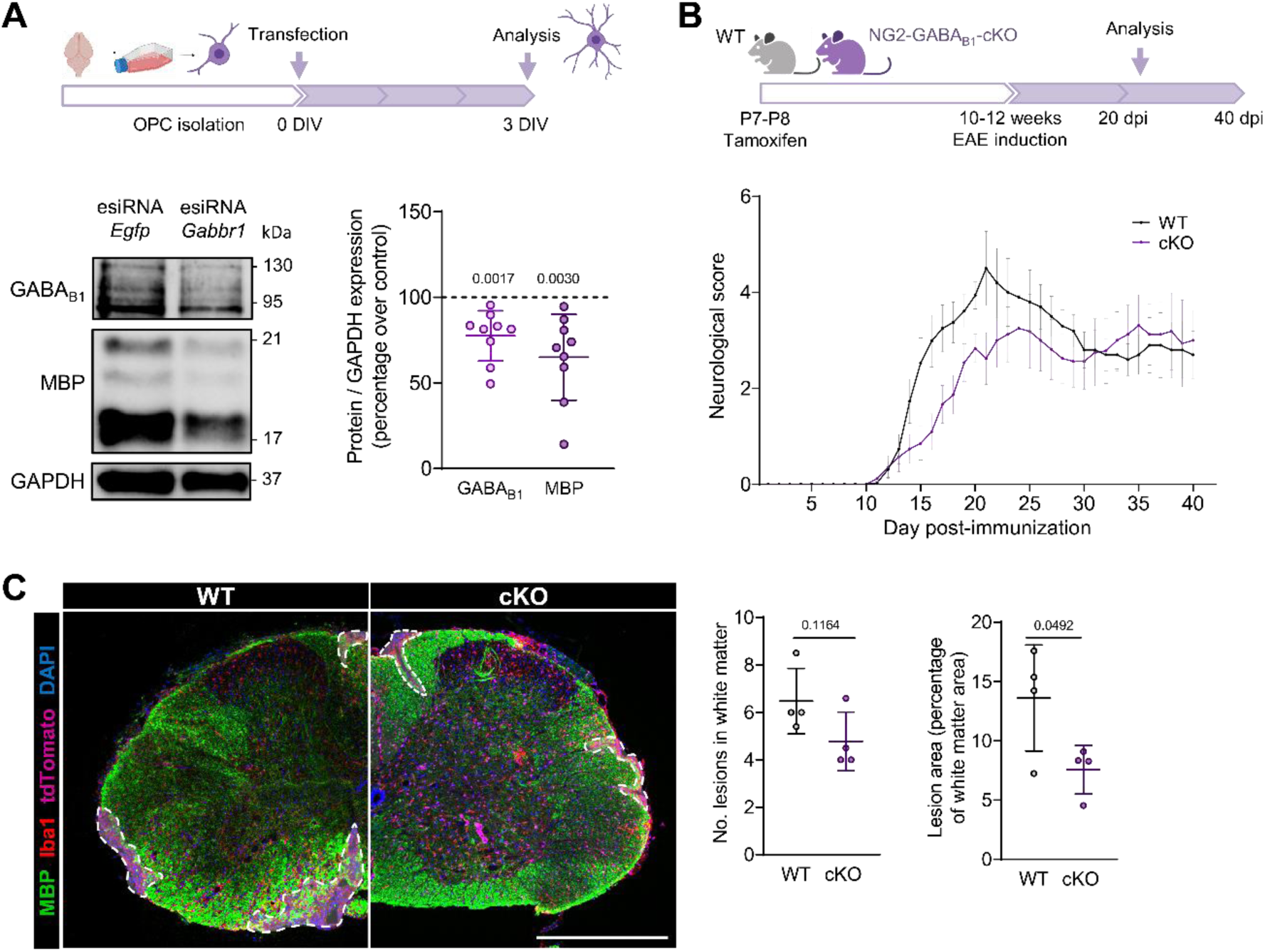
NG2-GABA_B1_ deletion reduces symptom severity in the acute phase of the EAE model. (A) Representative Western blot images and quantification of the expression of GABAB1 and MBP in 3 DIV cortical OPCs transfected with Egfp (control) or Gabbr1 esiRNA. Protein levels were normalized to GAPDH and expressed as percentage over control (cells transfected with Egfp; 100 % value). Paired one-way ANOVA. (B) Diagram showing the experimental setup for EAE induction in WT and cKO mice for GABA_B1_ in NG2^+^ cells and progression of the neurological score relative to motor symptoms in WT and cKO mice. Data represent mean ± SEM from at least 16 animals. (C) Representative confocal images presenting coronal sections of the spinal cord of WT and cKO mice in the EAE model at 20 dpi, immunostained with MBP (green), Iba1 (red) and DAPI (blue), tdTomato expression in the cKO is shown in magenta. White dashed lines indicate the lesions. Histograms show the number of demyelinating lesions and lesion size relative to white matter area. Unpaired Student’s t-test. Data represent mean ± SD and dots represent different animals or independent experiments. Scale bar: 500 μm.

### NG2-specific loss of GABA_B1_ drives immune gene expression in oligodendrocytes in acute EAE

Our silencing experiments, along with previous findings, showed that GABA_B_R blockade in isolated OPCs led to reduced MBP expression (Serrano-Regal et al., 2020), likely impairing their maturation. Thus, we next aimed to determine whether GABA_B1_ deletion in NG2^+^ cells affected oligodendrogenesis in the EAE model, and to evaluate if the observed reduction in the severity during the acute phase correlated with increased OPC differentiation. For that, we assessed the populations of OPCs (PDGFRα+Olig2+ cells), MOLs (APC+Olig2+ cells) and total oligodendrocytes (Olig2+ cells) by immunohistochemistry at 20 dpi in WT and cKO mice lesions and perilesions (Fig. 2a-c). Our analysis did not reveal any significant changes in the percentage of OPCs or MOLs present in lesions or perilesions. Moreover, no differences were found regarding the density of oligodendrocytes in these areas. Therefore, these data show that GABA_B1_ deletion in NG2+ cells does not modify OPC differentiation during the acute phase of the EAE.

**Figure 2.**
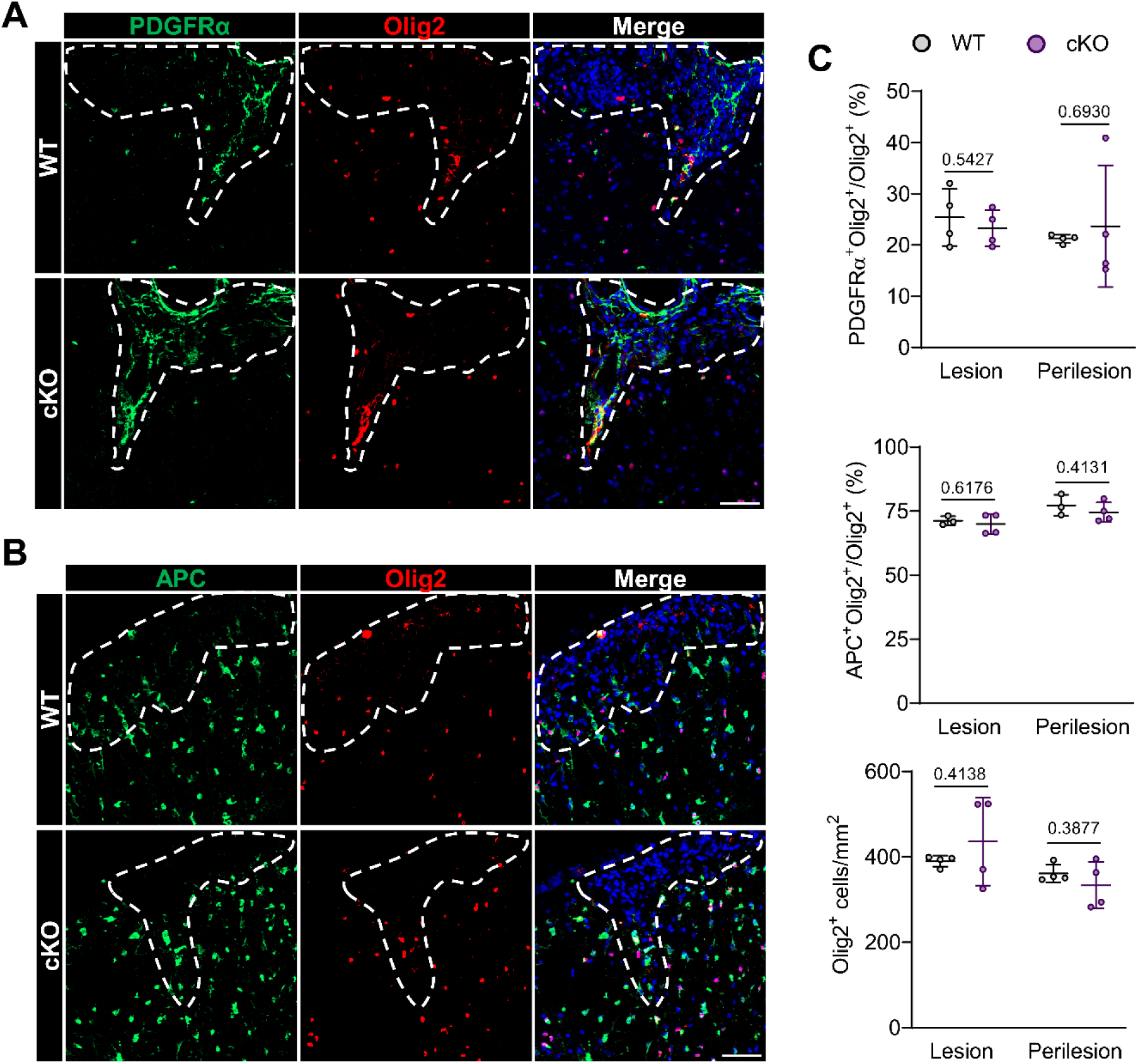
NG2-GABA_B1_ deletion does not modify oligodendrocyte differentiation in the acute phase of the EAE model. (A, B) Representative confocal images presenting coronal sections of the spinal cord of control and baclofen-treated mice in the EAE model at 20 dpi, immunostained with PDGFRα (green) (A) or APC (green) (B), Olig2 (red) and DAPI (blue). White-dashed lines indicate the lesion site. Scale bar: 50 μm. (C) Histograms showing the percentage of PDGFRα^+^Olig2^+^ cells from total Olig2^+^ cells, the percentage of APC^+^Olig2^+^ cells from total Olig2^+^ cells and the number of Olig2^+^ cells per area, in the lesion and perilesion (area near the lesion). Unpaired Student’s t-test. Data represent mean ± SD and dots represent different animals.

Next, to investigate the mechanisms underlying the protective effect of GABA_B1_ deletion in OPCs during the acute phase of the EAE, we analyzed molecular pathways potentially affected by this ablation. To achieve this and given the heterogeneity of oligodendrocyte populations in EAE (Falcão et al., 2018), we performed snRNA-seq of the spinal cord from WT and cKO mice subjected to EAE (Fig. 3a-c). Unsupervised clustering of the whole integrated dataset at low resolution revealed 9 distinct cell classes, based on their top differentially expressed genes (Fig. 3b). As GABA_B1_ deletion was specific for NG2^+^ cells, we next focused on the oligodendrocyte populations for more detailed analysis. Unsupervised clustering of the integrated OPC and MOL populations showed specific cell subtypes that were present in both WT and cKO. To identify potential differences induced by GABA_B1_ deletion during EAE in oligodendrocytes, we reprocessed both datasets without employing integration techniques to explore distribution in both WT and cKO samples across identified oligodendrocyte subtypes (Fig. 3c). This comparison showed similar distributions of both WT and cKO nuclei across the different oligodendrocyte subtypes. Intriguingly, these analyses revealed lower number of nuclei in the cKO sample compared to the WT. Nonetheless, despite cKO dataset presented less oligodendroglial nuclei, the percentage of nuclei conforming the subtypes was similar in most cases for both samples (Extended Fig. 1a).

**Figure 3.**
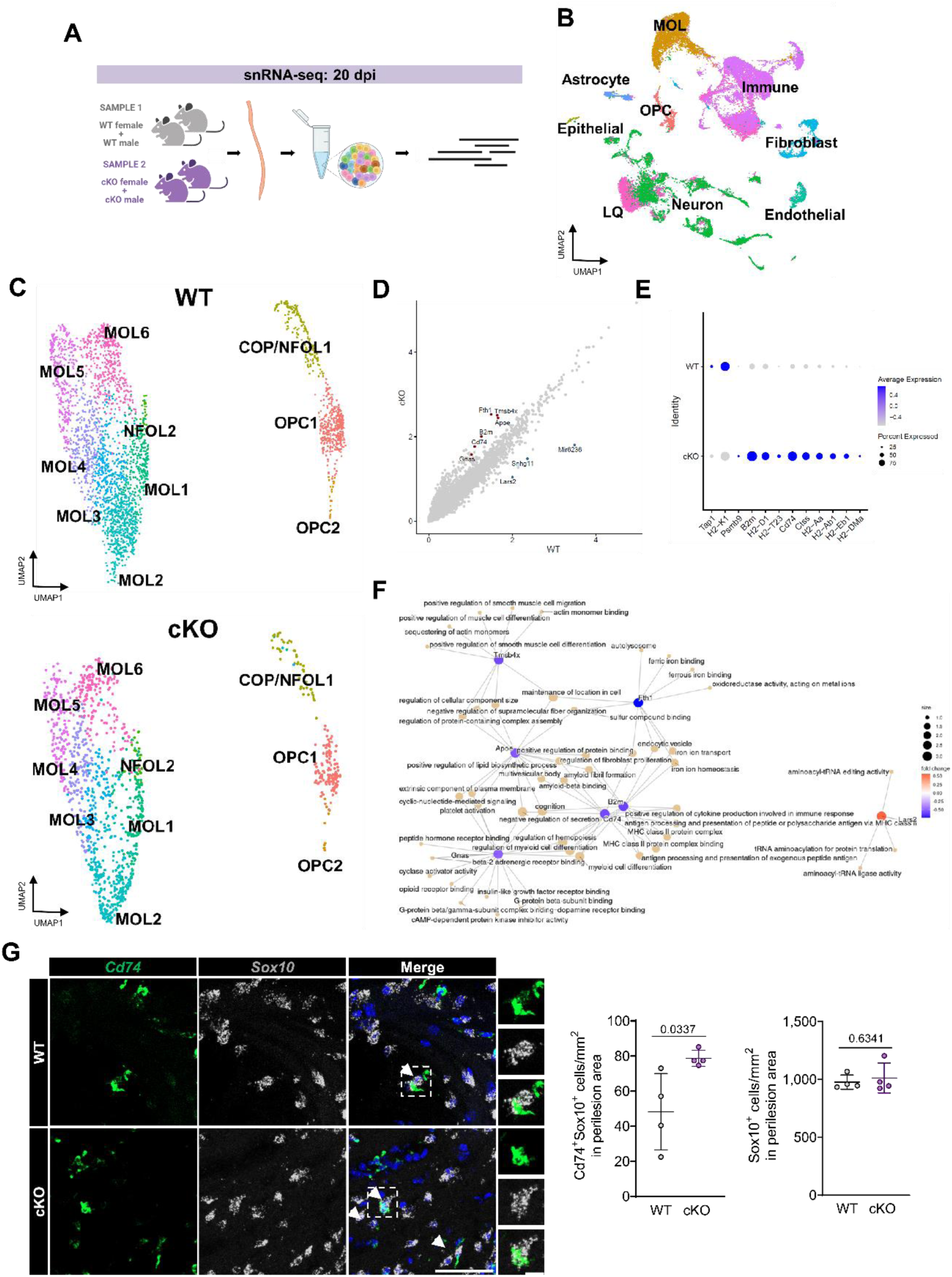
NG2-GABA_B1_ cKO shows upregulation of immune genes in oligodendroglia in the EAE model. (A) Diagram showing the experimental setup for sample processing of snRNA-Seq experiments. (B) Cell type annotation of unsupervised clustering of integrated WT and cKO samples. (C) Oligodendrocyte subtype annotation of unsupervised clustering of SCT WT (left) and cKO (right) samples. (D) Scatter plot showing differentially expressed genes in cKO mice compared to WT. Faint grey dots indicate no significant differences, blue dots indicate downregulation and red dots indicate upregulation. (E) Dot plot showing the expression pattern of immune genes in WT and cKO samples. (F) Over-representation analysis for the differentially expressed genes. (G) Representative confocal images presenting RNAscope *in situ* hybridization of coronal sections of the spinal cord of WT and cKO mice at 20 dpi, with probes for *Cd74* (green) and *Sox10* (grey) and stained with DAPI (blue). Histograms show the number of Cd74^+^Sox10^+^ and Sox10^+^ cells, Cd74^+^ cells in the perilesion area. Unpaired Student’s t-test. Data represent mean ± SD and dots represent different animals. Arrows point at Cd74^+^Sox10^+^ cells. Scale bar: 50 μm, higher magnification: 10 μm.

To identify candidate genes potentially influenced by GABA_B1_ deletion in this pathological context, we conducted differential gene expression analysis between WT and cKO in the whole oligodendrocyte population (Fig. 3d,f). We found 9 significantly differentially expressed genes between the two conditions. Specifically, *Tmsb4x*, *Fth1*, *Gnas*, *Apoe*, *B2m* and *Cd74* genes were upregulated in the cKO compared to WT, while *Mir6236*, *Snhg11* and *Lars2* genes were downregulated. When performing functional enrichment, immune functions were associated to upregulated genes in cKO oligodendroglia such as *B2m* and *Cd74*, members of members of major histocompatibility complex (MHC) I and II families, respectively. Histologically, *RNAscope* in situ hybridization at 20 dpi revealed a significant increase in the number of Cd74+Sox10+ per area in cKO mice (Fig. 3e). Besides, neither the oligodendroglial population, nor the Cd74+ population (data not shown) were altered in cKO mice compared to WT. Furthermore, although restricted differential gene expression analysis of the scRNA-Seq dataset did not reveal significant changes in the expression of *Gabbr1* in oligodendrocytes, we verified that this gene codifying GABA_B1_ subunit was downregulated in cKO mice compared to WT (Extended Fig. 1b). These results linked the cKO transcriptomic signature to an immune-associated phenotype, described in oligodendrocytes during EAE (Falcão et al., 2018; Meijer et al., 2022).

To further explore the link between GABA_B_R and the acquisition of an immune phenotype in oligodendrocytes, we treated rat cortical OPCs with IFNγ to mimic the inflammatory conditions characteristic of the EAE model. Then, we tested the impact of GABA_B_R activation with its agonist baclofen following insult with IFNγ. Although baclofen treatment did not seem to alter the increase in gene expression of *Cd74* or *Ciita* induced by IFNγ (Extended Fig. 2), analysis by immunocytochemistry showed that GABA_B_R activation with baclofen was able to revert the increase in MHC-II expression induced by IFNγ treatment in differentiating oligodendrocytes (Fig. 4a). Additionally, baclofen addition enhanced the number of mature oligodendrocytes both in the presence and absence of IFNγ. These observations provide evidence of the influence of GABA_B_R on the immune-related response of oligodendrocytes under inflammatory conditions. In addition, they advocate for an association between GABA_B_R deletion in OPCs during development and a potentially protective overexpression of immune-related genes during the EAE.

**Figure 4.**
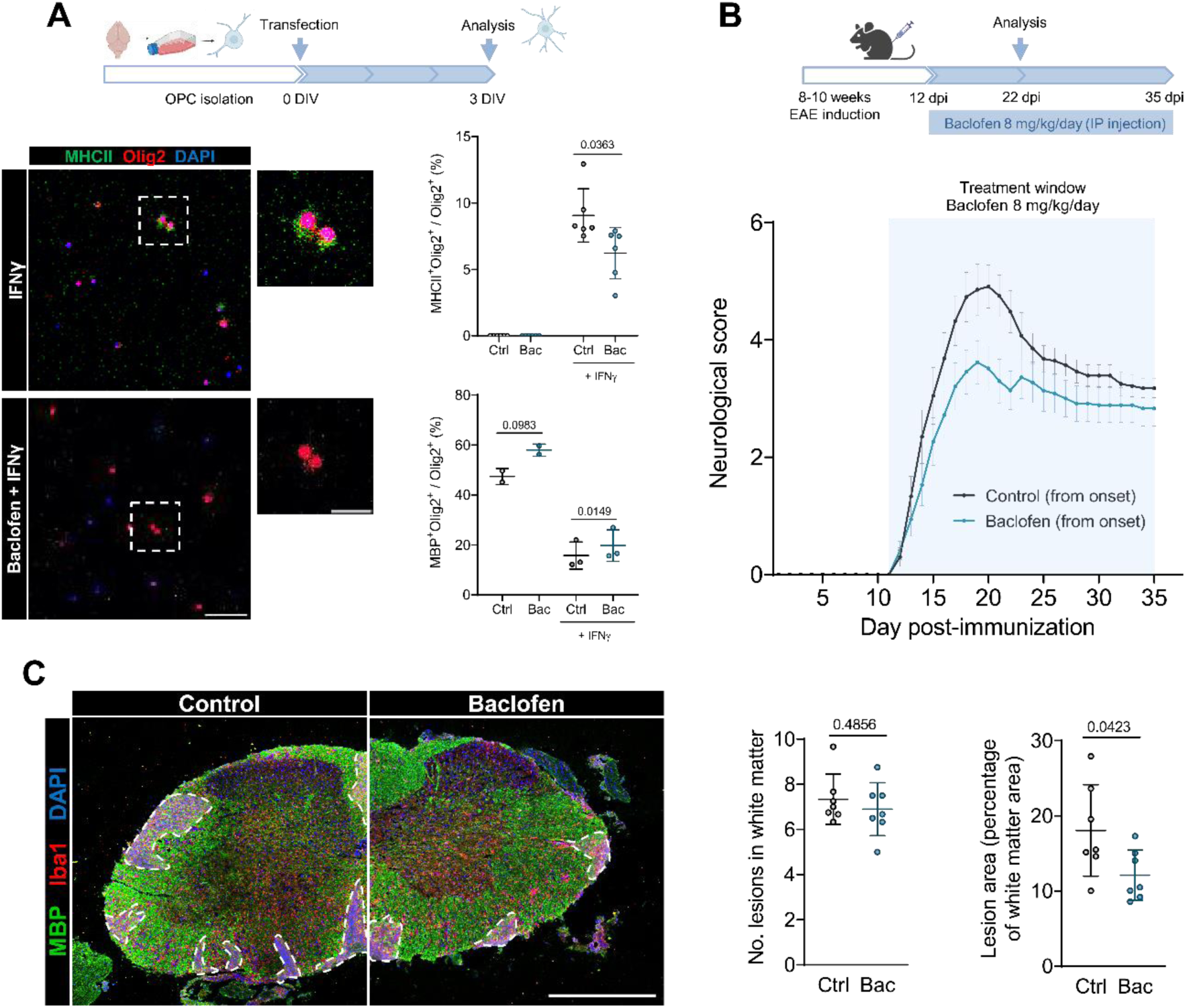
GABA_B_R activation with baclofen treatment ameliorates neurological symptoms and demyelinating lesion size in the EAE model. (A) Representative confocal images presenting rat cortical oligodendrocytes at 3 DIV treated with IFNγ and/or baclofen and immunostained with MHC-II (green), Olig2 (red) and DAPI (blue). Histograms show the percentage of MHC-II^+^ or MBP^+^ cells from total Olig2^+^ cells. Paired Student’s t-test. (B) Diagram showing the experimental setup for baclofen administration in the EAE model and progression of the neurological score relative to motor symptoms in control and baclofen-treated mice, with the treatment starting from the onset of the symptoms. Data represent mean ± SD from at least 14 animals. (C) Representative confocal images presenting coronal sections of the spinal cord of control and baclofen-treated mice in the EAE model at 22 dpi, immunostained with MBP (green), Iba1 (red) and DAPI (blue). White dashed lines indicate the lesions. Histograms show the number of demyelinating lesions and lesion size relative to white matter area. Unpaired Student’s t-test. Data represent mean ± SD and dots represent different animals. Scale bars: A = 25 μm, D = 500 μm; C = 50 μm.

Therapeutic baclofen promotes OPC differentiation and reduces T cell entry, alleviating EAE Furthermore, to assess the therapeutic potential of baclofen in a complex MS model, we investigated its effects on EAE progression. Hence, we induced EAE in young adult mice and administered baclofen daily via IP injection from 12 dpi until analysis (Fig. 4b). In this case, our treatment window started at the onset of the symptoms, aligning with a potential therapeutic approach. Remarkably, our initial injections revealed that baclofen treatment induced a transient state of lethargy in the mice during the first hour following the injection, presumably due to the systemic activation of inhibitory neurotransmission induced by the drug. However, afterwards the animals were fully awake and displayed apparent normal behavior, similar to control mice. Daily monitoring of the neurological symptoms also revealed a reduction in the severity of the EAE course in baclofen-treated mice compared to controls, with no differences in the onset of the symptoms. Interestingly, we also examined the effect of GABA_B_R blockade on the course of the EAE, administering GABA_B_R antagonist CGP35348 via IP injection from 12 dpi (Extended Fig. 3). In this case, no remarkable differences were observed in the neurological score of the mice, except for a slight deterioration during the recovery phase. Immunofluorescence analysis of demyelinating lesions at 22 dpi following baclofen treatment revealed a decrease in the percentage of the white matter area affected by lesions, paradoxically similar to GABA_B1_ deletion (Fig. 4c). In summary, these findings, while indicating that modulation of GABA_B_R is likely affecting several pathways that have opposite effects during neuroinflammation, also emphasize the potential of baclofen as a promising agent for remyelinating therapeutic strategies against MS, providing evidence of its effect in a model that closely mirrors the full spectrum of the characteristics of this disease.

The critical role of OPC differentiation for remyelination and recovery within this pathological context led us to investigate the impact of baclofen treatment on oligodendroglia in EAE demyelinating lesions. With this aim, we conducted immunofluorescence analyses to evaluate the presence of OPCs and mature oligodendrocytes relative to the total oligodendroglial population at 22 dpi (Fig. 5a, b), both within the lesion and perilesion areas. We found a significant decrease in the percentage of OPCs among the total number of oligodendrocytes in the lesions, as well as a non-significant reduction in the perilesions. In this line, we verified that baclofen administration provoked a significant increase in the percentage of MOLs relative to the total oligodendroglial population, both within the lesion and perilesion area. Notably, we did not find any modifications in the total number of oligodendrocytes per area. These observations demonstrate that baclofen treatment boosts OPC differentiation within demyelinating lesions in the EAE model.

**Figure 5.**
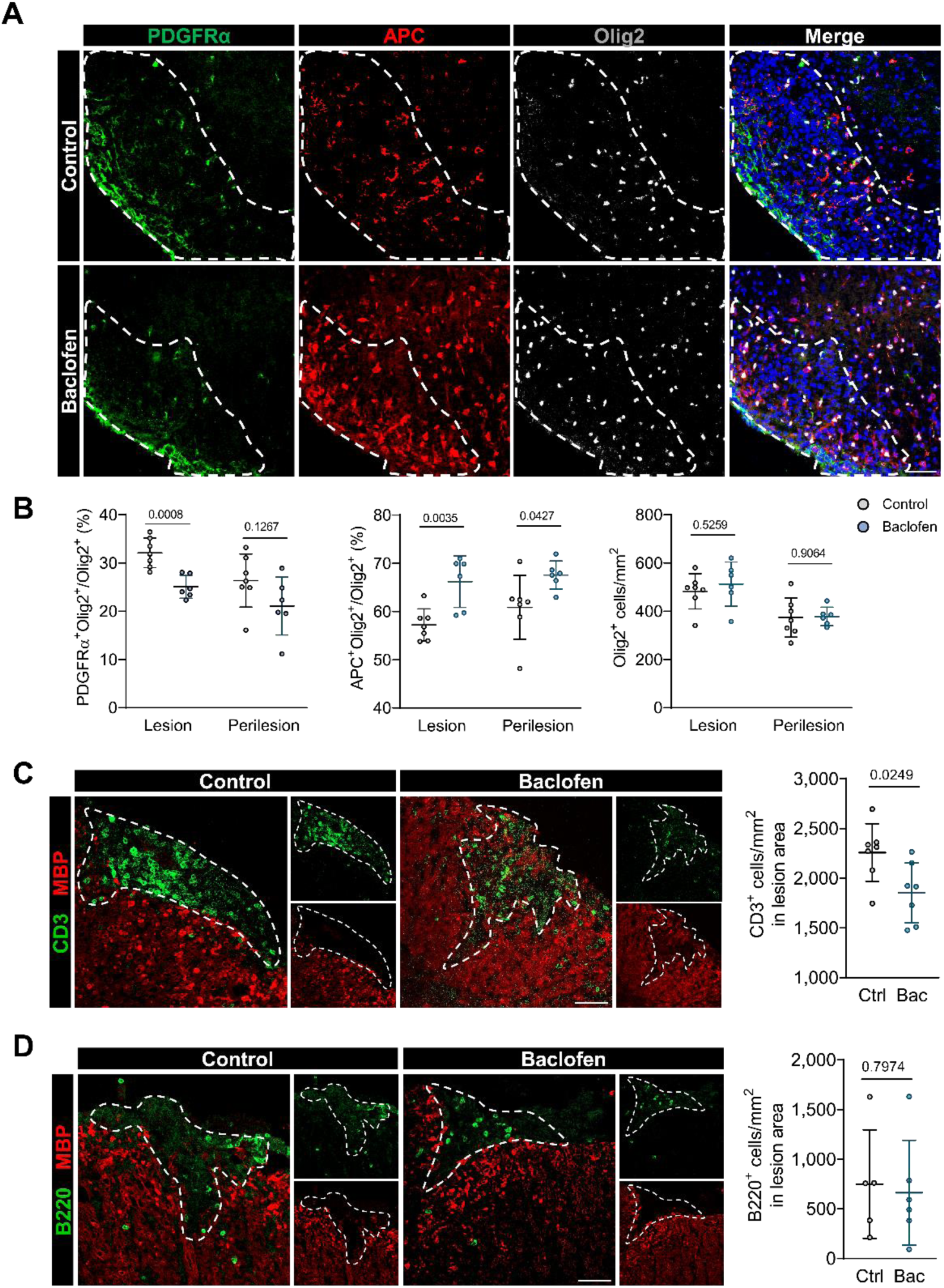
Baclofen-mediated GABA_B_R activation enhances OPC differentiation and limits T cell infiltration in EAE lesions. (A) Representative confocal images presenting coronal sections of the spinal cord of control and baclofen-treated mice in the EAE model at 22 dpi, immunostained with PDGFRα (green), APC (red), Olig2 (gray) and DAPI (blue). White-dashed lines indicate the lesion site. (B) Histograms showing the percentage of PDGFRα^+^Olig2^+^ cells from total Olig2^+^ cells, the percentage of APC^+^Olig2^+^ cells from total Olig2^+^ cells and the number of Olig2^+^ cells per area, in the lesion and perilesion. (C, D) Representative confocal images presenting coronal sections of the spinal cord of control and baclofen-treated mice in the EAE model at 22 dpi, immunostained with CD3 (green) (C) or B220 (green) (D) and MBP (red). White-dashed lines indicate the lesion site. Histograms show the number of CD3^+^ (C) or B220^+^ (D) cells per lesion area. Unpaired Student’s t-test. Data represent mean ± SD and dots represent different animals. Scale bar: 50 μm.

The interaction between cells from the immune system and the CNS is key in the development and pathogenesis of EAE. T cells are primarily responsible for the infiltration into the CNS, with B cells also contributing to this process, and subsequently mediating the response that leads to demyelination and damage (Constantinescu et al., 2011). Thus, we examined the presence of both T cell (CD3^+^ cells) and B cell (B220^+^ cells) populations in the lesions of control and baclofen-treated mice at 22 dpi using immunofluorescence. We detected a significant reduction in the number of T cells per lesion area following baclofen administration (Fig. 5c), whereas no alterations were observed in the density of B cells per lesion area (Fig. 5d). These findings indicate that baclofen administration may alter T cell infiltration in EAE lesions. Overall, our data indicates that GABA_B_R signaling is multifactorial, acting in distinct cellular contexts to shape disease progression. Modulating GABA_B_R signaling at different stages of disease evolution may therefore produce divergent outcomes.

### Prophylactic baclofen delays EAE onset by modulating immune responses

Given its impact on the T cell population and the critical role of T cell priming and infiltration in this model (Constantinescu et al., 2011), we next investigated whether baclofen treatment during EAE development could influence disease progression. To address this, we induced EAE in young adult mice and administered baclofen starting at the day of immunization. Notably, we found that the treatment resulted in a marked delay in the onset of neurological symptoms while still reducing their severity, albeit to a lesser extent than in the previous experimental setup (Fig. 6a). These findings suggest that baclofen may modulate T cell populations, highlighting its potential immunomodulatory role in addition to its demonstrated pro-differentiating effects on oligodendroglia.

**Figure 6.**
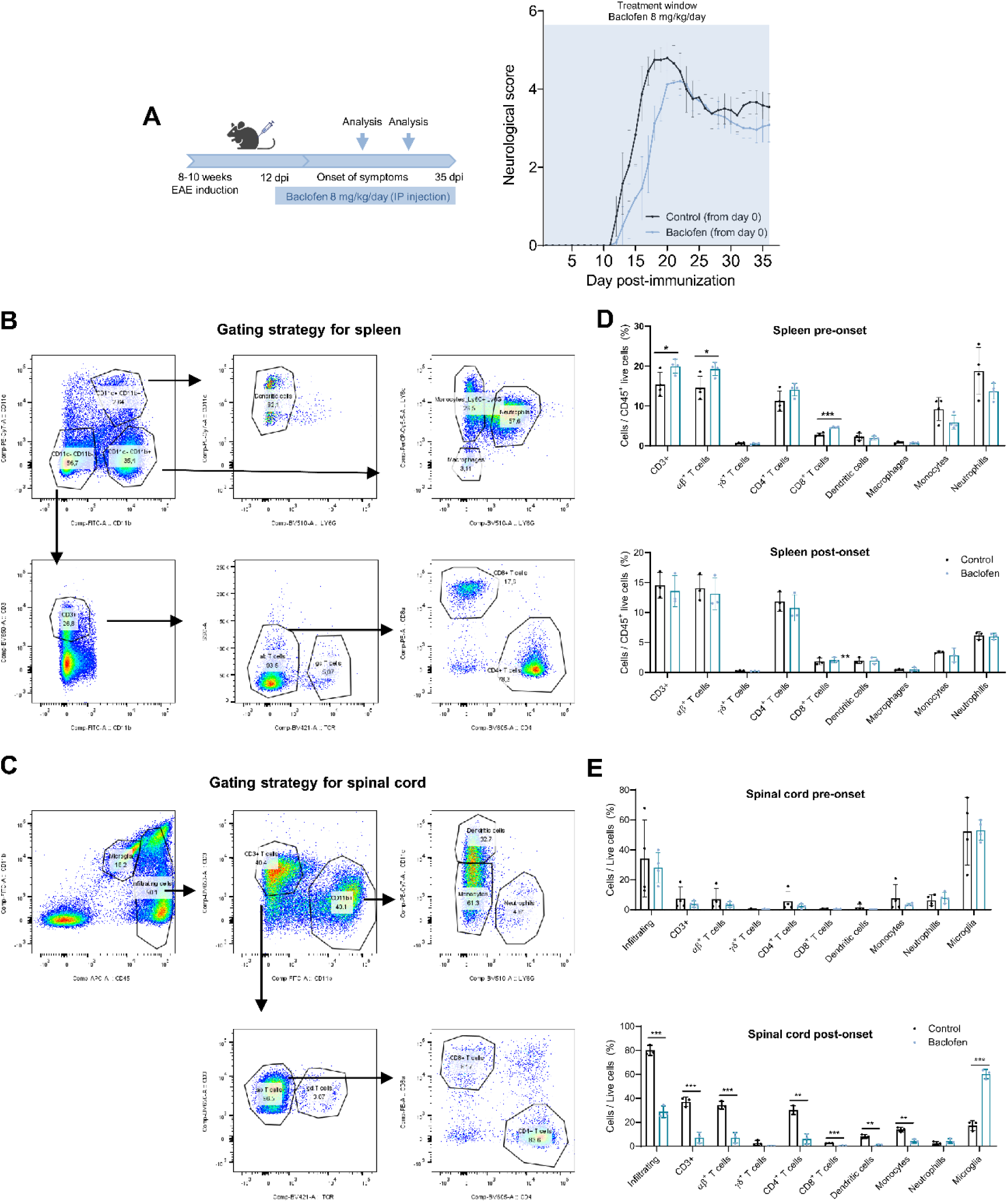
GABA_B_R activation with baclofen affects immune cell activation in the EAE model. (A) Diagram showing the experimental setup for baclofen administration in the EAE model and progression of the neurological score relative to motor symptoms in control and baclofen-treated mice, with the treatment starting from the start of the model. Data represent mean ± SD from 6 animals. (B, C) Flow cytometry gating strategy for analysis of immune populations from the spleen (B) or spinal cord (C) of mice at EAE pre-onset. (D, E) Histograms showing the percentage of immune cell populations present in the spleen (D) or spinal cord (E) at EAE pre-onset or post-onset. Unpaired Student’s t-test. Data represent mean ± SD and dots represent different animals.

To further investigate the impact of baclofen treatment on immune cell activation and clarify the link between GABA_B_R and immune cell responses in the EAE model, we conducted flow cytometry analysis to assess changes in immune cell populations in the spleen and spinal cord of control and baclofen-treated EAE mice (Fig. 6b, c). Prior to symptom onset, our data showed an increase in the presence of αβ^+^ and CD8^+^ T cells in the spleen of mice treated with baclofen, whereas no significant alterations were observed after symptoms appeared (Fig. 6d). In contrast, analysis of the spinal cord revealed a pronounced reduction in major immune cell populations following treatment with baclofen during the EAE (Fig. 6e). Notably, unlike other immune cells, the microglia population was significantly increased in the spinal cord of baclofen-treated animals. These results indicate that GABA_B_R activation reduces immune cell infiltration in the spinal cord and might influence immune cell activation, further demonstrating its importance to modulate immune activity and highlighting its role in modulating immune responses beyond its effects on oligodendrocyte differentiation.

## Discussion

Remyelination remains a major unmet challenge in current MS therapies, in a landscape with an increasing need for strategies to address the progressive stages of the disease. Yet, remyelination alone may be insufficient to address the complex pathophysiology of MS. Effective treatments should combine immunomodulation—to prevent new inflammatory relapses—with approaches that promote remyelination, repair existing damage, and provide neuroprotection. In this context, we identify GABA_B_ R as a promising dual-action target, capable of regulating both oligodendroglial differentiation and immune responses across multiple cell types.

Recent research points at GABA_B_R as a relevant modulator of oligodendrocyte functionality (Serrano-Regal et al., 2020b; Bai et al., 2021). In this context, our initial finding demonstrating that GABA_B1_ silencing impairs MBP expression in cultured OPCs align with previous observations showing that GABA_B_R blockade using its antagonist CGP55845 reduced OPC differentiation (Serrano-Regal et al., 2020). These results further support the constitutive role of this receptor in oligodendrocyte maturation. Nonetheless, the milder phenotype observed during the acute phase of the EAE in NG2-GABA_B1_ cKO mice suggests either a novel role of GABA_B_R activity when its manipulation was restricted to OPCs, or an effect specific to the inflammatory context of EAE. Although increased oligodendrocyte differentiation was observed in the lysolecithin and cuprizone models when deleting GABA_B1_ in NG2+ cells at 10 weeks of age (Gobbo et al., 2024), NG2-GABA_B1_ deletion at P7-9 is known to lead to hypomyelination of interneurons and reduced OPC maturation during development (Fang et al., 2022). Moreover, deletion of GABA_B1_ in Schwann cells has also been shown to compromise myelination (Faroni et al., 2019), all supporting the essential role of GABA_B1_ in OPC differentiation and myelination under physiological conditions. However, our findings suggest that GABA_B_R function may be altered under inflammatory and pathological conditions, characteristic of the EAE model.

Notably, the absence of alterations in oligodendrocyte maturation state in our NG2-GABA_B1_ cKO mice suggests that the differences observed during the acute phase of the EAE are not primarily driven by major changes in the differentiation process, but rather by alternative mechanisms. While we cannot exclude the potential contribution of other differentially expressed genes to the observed phenotype, we were particularly intrigued by the immune related genes *B2m* and *Cd74* identified in the transcriptomic analysis, which were enriched in oligodendrocytes from NG2-GABA_B1_ cKO mice. *B2m* and *Cd74* encode components from the MHC class I and II, respectively, and according to the snRNA-seq data, they are consistently upregulated in the cKO following EAE induction. Recent studies have described that oligodendrocytes can acquire an immune-like phenotype during EAE (Zeis et al., 2016; Falcão et al., 2018; Kirby et al., 2019; Meijer et al., 2022), indicating a potential role for these cells as modulators of the progression of this model. In our study, we verified that the small proportion of Cd74+ oligodendrocytes was increased in the cKO mice compared to the WT control. These findings indicate that deletion of GABA_B1_ in OPCs promotes the emergence of an immune phenotype in oligodendrocytes.

In fact, the acquisition of an immune phenotype by oligodendrocytes does not necessarily correlate with a worse outcome. While the functional relevance of these disease-associated oligodendroglia in MS remains unclear, they have been proposed to act as a brake on escalating neuroinflammation (Castelo-Branco et al., 2024). Although it seems clear that external inflammatory stimuli impair OPC differentiation (Ahn et al., 2021; Chew et al., 2005), the upregulation of the immune phenotype in oligodendrocytes might promote their own regenerative capacity through autocrine mechanisms, as observed in the cuprizone model (Moyon et al., 2015). Additionally, the bidirectional communication between oligodendrocytes and immune cells can modulate T cell responses (Cabeza-Fernández et al., 2023), which in our study may contribute to the attenuation of the acute phase in the EAE. Moreover, oligodendrocytes expressing MHC class II genes have been reported to phagocytose myelin (Falcão et al., 2018). Considering the critical role of microglia in clearing myelin debris to favor neuroprotection (Cignarella et al., 2020; Sen et al., 2022), it is plausible that enhanced immune functionality in oligodendrocytes, resulting from GABA_B1_ ablation, exerts a protective effect. This mechanism could contribute to the reduced severity observed during the acute phase of the EAE. Taken together, these hypotheses reveal previously unrecognized mechanisms by which GABA_B_R signaling could modulate oligodendrocyte function. Next, with the aim of verifying the effect of GABA_B_R modulation in oligodendrocytes, we treated isolated oligodendrocytes with GABA_B_R agonist baclofen. We found a decrease in MHC-II expression in response to baclofen in IFNγ-treated oligodendrocytes, supporting a role for GABA_B_R in modulating oligodendrocyte immune responses under inflammatory conditions.

Given the decrease in oligodendrocyte immune phenotype promoted by baclofen, we next explored its impact in the EAE model. We had previously demonstrated the protective effect of baclofen in the lysolecithin model (Serrano-Regal et al., 2022). However, the EAE model is based on the autoimmune etiology attributed to MS, and more accurately replicates the main pathological features of the disease, including inflammation, demyelination, axonal loss and gliosis (Constantinescu et al., 2011; Glatigny & Bettelli, 2018). This provides a more complex environment that closely resembles the pathophysiology of MS. Furthermore, given the results observed in the NG2-GABA_B1_ cKO in EAE, we considered highly relevant to explore the effect of baclofen administration in this context.

Remarkably, baclofen treatment started at the onset of symptoms attenuated EAE severity, reducing the generation of demyelinating lesions within the spinal cord white matter. Starting the treatment at the symptom onset not only increased the therapeutical relevance of our approach but also minimised interference with the early inflammatory mechanisms that guide EAE development. This finding reveals a novel role for GABA_B_R activation by baclofen when administered from the onset of the symptoms.

Previous studies have reported no significant changes in EAE progression following baclofen treatment, although in those cases baclofen was administered after the peak of the symptoms (Ghareghani et al., 2018). In line with our observations, enhanced GABAergic activity has been reported to ameliorate symptomatology in EAE (Bhat et al., 2010; Stamoula et al., 2023; Tian et al., 2018), further highlighting the relevance of GABA receptors for MS pathology.

The improved outcome in EAE following baclofen treatment was most likely not related to a reduction in oligodendroglia with immune properties, but rather was accompanied by increased OPC differentiation within lesion and perilesion areas, consistent with our previous findings in the LPC model (Serrano-Regal et al., 2022) and from studies in isolated OPCs (Serrano-Regal et al., 2020). Since no differences in oligodendrocyte maturation were observed during EAE in the NG2-GABA_B1_ cKO, we hypothesize that GABA_B_R activation in other cell types such as GABAergic interneurons, and not only in oligodendrocytes may contribute to the observed increase in oligodendrocyte differentiation in this context. Indeed, baclofen treatment has been shown to increase the number of OPC-interneuron pairs present in the mouse cortex, potentially promoting bidirectional communication (Boulanger & Messier, 2017) and enhancing oligodendrocyte lineage progression.

In addition, we found reduced T cell infiltration in EAE lesions following baclofen administration at day 12, suggesting that baclofen might attenuate the inflammation relative to this model. To further explore this effect, we started baclofen treatment from day 0 and monitored immune cell density in both the spleen and spinal cord before and after symptom onset. Interestingly, while several immune cell populations were increased in the spleen of baclofen-treated mice prior to the appearance of the first symptoms, a strong reduction in immune cell infiltration was detected in the spinal cord. These findings are in line with previous studies showing that modulation of GABAergic signaling can regulate the autoimmune response in EAE (Stamoula et al., 2023), and even that baclofen treatment can reduce inflammation in other disease models (Duthey et al., 2010; Huang et al., 2015). Furthermore, these alterations in immune cell activation could also be a consequence of baclofen-induced effects in oligodendrocytes, as these have been shown to modulate immune response (Wang et al., 2025). It is possible that reduction of immune gene expression in oligodendroglia by baclofen, while affecting their potential protective role, could lead to reduced recruitment of T cells from the periphery. Notably, microglia population was the only cell type in which we observed an increase following baclofen administration in the mouse spinal cord. Microglia are highly versatile cells capable of adopting diverse functional states beyond the classical anti- or pro-inflammatory phenotypes (Paolicelli et al., 2022) and they are known to be key contributors to remyelination (Miron et al., 2013; Lloyd et al., 2019). Thus, we propose that baclofen can also target microglia during the early phases of EAE, contributing to the protective effect exerted by the drug.

Based on the results described in this study and considering the established use of baclofen in managing spasticity in MS, we propose that this compound is a promising candidate for drug repurposing. It could not only act as a pro-remyelinating drug, but we have now demonstrated that GABA_B_R modulation with baclofen is a relevant immunomodulatory approach. Although its efficacy in promoting remyelination in patients remains to be fully evaluated, earlier administration of this drug could potentially enhance myelin repair. However, the transient lethargic state observed in mice following baclofen administration raised awareness about the safety and potential side effects of the drug. These sedative effects have been previously reported with high intraperitoneal doses of baclofen (Li et al., 2013). Nevertheless, the recommended dose for patients is 80 mg/day (Kent et al., 2020), which is lower than the dose used in our study, despite similar high doses being employed in rodent models (Boulanger & Messier, 2017; Ghareghani et al., 2018). High doses of baclofen can lead to adverse outcomes, including toxicity and withdrawal syndrome, although such complications are more frequently associated with intrathecal administration (Romito et al., 2021). Therefore, although no additional side effects were detected in the animals used in this study, the safety profile of baclofen should be carefully considered to ensure safe clinical use.

While this study provides novel insights into the role of GABA_B_R in oligodendrocytes, future research will address current limitations, including the effects of baclofen impact on other cell types such as immune cells and microglia, as off-target actions may contribute to the observed outcome. Moreover, the analysis of samples from patients with MS treated with baclofen will help determine its clinical potential in MS, taking into account personalized treatment windows and patient-specific inflammatory contexts.

## Materials and methods

### Animals

All experiments with animals were reviewed and approved by the internal Animal Ethics Committee of the University of the Basque Country (UPV/EHU), according to the guidelines set by the European Communities Council Directive 2010/63/EU. Protocols were evaluated and sanctioned by the Ethics Committee on Animal Experimentation (CEEA), the collegiate authority that operates under the Ethics Commission for Research and Teaching (CEID) at the UPV/EHU. Animals were housed under standard conditions, maintaining a 12 h light/dark cycle and ad libitum access to food and water. Every possible measure was taken to minimize animal suffering and the number of animals used per experiment.

Experiments to evaluate GABA_B1_ deletion in OPCs in the EAE model were conducted on 10-12 week old female and male WT and cKO mice for GABA_B1_ deletion, with cKO mice expressing tdTomato in NG2+ cells/OPCs (NG2-CreERT2:GABA_B1_ ^fl/fl^;tdTomato mice or NG2-GABA_B1_ cKO, from the laboratory of Dr. Frank Kirchhoff). For the GABA_B1_ transgenic line, recombination was induced by two intraperitoneal (IP) tamoxifen injections (100 mg/kg) at postnatal day 7 and 8. EAE experiments with baclofen administration were performed using 8-10 week old C57BL/6J female mice. For primary cell cultures, male and female P0-2 Sprague Dawley rats were used.

### Experimental autoimmune encephalomyelitis

For the assessment of a model mimicking several MS features, EAE was induced in 8 to 10-week-old C57BL/6J female mice or in female and male WT and cKO mice for NG2-GABA_B1_. Mice were immunized via three subcutaneous injections in the hindlimbs with MOG_35-55_ (66.67 µg per injection), with the peptide sequence MEVGWYRPFSRVVHLYRNGK. MOG_35-55_ was emulsified in incomplete Freund’s adjuvant (Sigma-Aldrich, #F5506) supplemented with 8 mg/mL of *Mycobacterium tuberculosis* H37Ra (BD #231141). Pertussis toxin (Sigma-Aldrich #516560) diluted in water was administered via IP injection (500 ng per day) on the day of immunization and 2 days later to ensure model development. Animals in this model exhibit progressive motor symptoms, which were daily monitored and scored on a scale from 0 to 8 as follows: 0, no detectable changes in motor behavior; 1, weakness of the tail; 2, paralyzed tail; 3, impairment or weakness in hindlimbs; 4, hemiparalysis of hindlimbs; 5, complete hindlimb paralysis; 6, hindlimb paralysis and rigidity in forelimbs; 7, tetraplegia; 8, moribund.

For baclofen treatment during the EAE model, mice were administered daily IP injections of saline solution or baclofen (8 mg/kg) either from the day of immunization (0 dpi) or the onset of the symptoms (12 dpi). Animals were then sacrificed and processed in accordance with the subsequent experimental procedure.

### Oligodendrocyte primary cell culture

Primary mixed glial cultures were obtained from the cortical lobes of P0-2 Sprague Dawley rats, according to previously described protocols (McCarthy & de Vellis, 1980) with some modifications (Sánchez-Gómez et al., 2018). Forebrains were removed and dissected to isolate cortices and these were then subjected to enzymatic and mechanical digestions. The resulting cell suspension was in 75 cm^2^ flasks previously coated with 1 µg/mL poly-D-lysine (PDL; Sigma-Aldrich #P0899) diluted in water. Flasks were maintained in culture conditions at 37 °C with 5 % CO2 in Iscove’s modified Dulbecco’s medium (IMDM, Gibco #42200-014) supplemented with 10 % Hyclone fetal bovine serum (FBS Hyclone; Gibco #SH30071.03IH30-45). This glial culture contained a mixture of OPCs, microglia and astrocytes. For OPC isolation, flasks were first subjected to mechanical shaking after 7 or 14 days in culture to detach microglia. The remaining OPCs within the astrocyte monolayer were separated by overnight shaking and filtering through 10 μm pore size nylon meshes. OPCs were seeded onto PDL-coated coverslips placed in 24-well plates at 10,000 cells/well for analysis by immunocytochemistry and at 60,000 cells/well for Western blot.

For drug treatments, OPCs were treated with of IFNγ (100 ng/mL) on the seeding day (day 0), and cells were processed 72 h later, on day 3. GABA_B_R agonist baclofen (100 μM; Tocris #0796) was added daily. In treatments combining baclofen with IFNγ, on day 0 GABAergic drugs were added in the first place and incubated for 30 min before the addition of IFNγ.

For gene silencing assays, OPCs were transfected with 8 µM of MISSION esiRNA targeting *Egfp* or *Gabbr1* (Sigma-Aldrich) using the Amaxa Basic Nucleofector Kit for Primary Mammalian Glial cells (Lonza #VPI-1006), following the manufacturer’s instructions. Briefly, at least 1,000,000 cells were resuspended in Amaxa solution containing the esiRNA and they were transfected using Amaxa Nucleofector 1 electroporation system (O-17 program). Subsequently, cells were resuspended in Opti-MEM medium (Gibco #31985070) and seeded as previously described. After one hour, Opti-MEM medium was replaced with SATO + medium and cells were processed for Western blot analysis at day 3.

### Immunofluorescence

For the immunofluorescence analysis of spinal cord tissue, mice were perfused with PBS for 2 min and spinal cords were post-fixed overnight at 4 °C in 4 % PFA diluted in PBS. Then, spinal cords were cryoprotected in 15 % sucrose (w/v) diluted in PBS for 48-72 h. After that time, the lumbar area of the spinal cord was cut into four 2-mm-thick blocks and placed in 15 % sucrose-7 % gelatin (w/v) diluted in PBS, and then frozen in isopentane for 2 min at -65 °C. Tissue was stored at -80 °C for appropriate tissue preservation. Coronal 12 μm-thick spinal cord sections were cut into Superfrost glass slides (Thermo Fisher #11976299) using a CM3050 S cryostat (Leica Biosystems), and stored at -20 °C until further experimentation. For immunocytochemistry, cells were fixed in the same 4 % PFA solution for 15 min at RT and washed for 3 times in PBS for 10 min at RT.

To perform immunohistochemistry, spinal cord sections were air-dried for 1 h at RT and rehydrated in PBS for 30 min. For the use of primary antibody against Olig2, antigen retrieval was carried out by adding low-pH R-Universal retrieval buffer (Aptum Biologics #AP0530-500) and heating the slices in a microwave for 45 s. For experiments using primary antibodies against MBP, slices were previously permeabilized in absolute ethanol for 10 min at -20 °C, followed by three washes in PBS for 10 min at RT. Tissue sections or coverslips containing cells were incubated for 30 min at RT in blocking solution containing 4 % NGS and 0.1 % triton X-100 diluted in PBS. Next, primary antibodies were added overnight at 4 °C in antibody solution containing 1 % NGS and 0.1 % triton X-100 diluted in PBS. Primary antibodies were: APC (1:200, Calbiochem #OP80), B220/CD45R (1:200, BD Biosciences #557390), CD3 (1:50, Bio-Rad #MCA1477), Iba1 (1:300, Synaptic systems #234004), MBP (1:1000, Biolegend #808402), MHC-II (1:300, abcam #ab23990), Olig2 (1:500, Millipore #MABN50), PDGFRα (1:500, R&D Systems #AF1062). The next day, sections were washed three times in PBS and incubated for 1 h at RT with Invitrogen AlexaFluor-conjugated secondary antibodies at a 1:500 dilution and DAPI (4 μg/mL) diluted in antibody solution. Secondary antibodies were: IgG Goat 488 (#A11055), IgG Guinea Pig 647 (#A21235), IgG2a Mouse 594 (#A21131), IgG2a Mouse 647 (#A21241), IgG2b Mouse 488 (#A21141), IgG2b Mouse 594 (#A21145), IgG Rabbit 488 (#A11008), IgG Rat 488 (#A11006). Finally, sections were additionally washed three times with PBS ant mounted with ProLong or Glycergel (Dako #C056330-2) mounting medium using coverslips for microscopy analysis.

### RNAscope *in situ* hybridization

Mice were perfused with 4 % PFA diluted in PBS and spinal cords were collected and snap frozen in liquid nitrogen, before further processing for cryostat cutting as previously described. The RNAscope protocol was applied using the RNAscope Multiplex Fluorescent Assay v2 (ACD #323110), according to the manufacturer’s instructions.

First, tissue sections were incubated for 4 min at 98 °C in target retrieval solution (RNAscope Target Retrieval Reagents, ACD #322000), and washed twice in DNase/RNase-free water for 2 min at RT. Afterwards, sections were incubated for 20 min at RT with protease IV (RNAscope Protease III & Protease IV Reagents, ACD #322340) and washed twice in DNase/RNase-free water. The tissue was then incubated for 2 h at 40 °C with specific probes for Sox10 (ready to use, ACD #435931) in C1 channel and Cd74 (1:50, ACD #437501) in C2 channel to ensure hybridization, followed by two washes using wash buffer (RNAscope Wash Buffer Reagents, ACD #310091). The hybridization signal was amplified by incubation at 40 °C with v2Amp1 (30 min), v2Amp2 (30 min) and v2Amp3 (15 min) (RNAscope Multiplex Fluorescent Detection Reagents v2, ACD #323110), with two washes using wash buffer between incubations. Subsequently, spinal cord sections were blocked by incubation for 15 min at 40 °C with v2-HRP-C1, followed by two washes with wash buffer. Then, they were incubated for 30 min at 40 °C with TSA-conjugated fluorophores (1:5000) diluted in TSA buffer (RNAscope Multiplex TSA Buffer, ACD #322810) to generate the fluorescent signal associated with the C1 channel. After two additional washes with wash buffer, similar incubations for blocking with v2-HRP and TSA-conjugated fluorophores were repeated for the C2 channel. Specifically, fluorescein (Perkin Efmer #FP1168015UG) and cyanine 5 amplification reagents (Perfin Efmer #FP1171801) were used as TSA-conjugated fluorophores. Finally, sections were incubated for 5 min at RT with DAPI (4 μg/mL), washed with PBS and mounted onto glass slides with ProLong mounting medium for visualization.

### Image acquisition and analysis

Image acquisition was conducted on one of the following confocal microscopes: Zeiss LSM800, Zeiss LSM880 Airyscan (SGiker, UPV/EHU) or Leica TCS SP8 (Achucarro Basque Center for Neurosciences). Same settings were kept within each experiment. 3D Histech Panoramic MIDI II slide scanner was used for the overview of EAE mouse spinal cord tissue. For images obtained using the slide scanner, annotations of the areas of interest were performed in each tissue section with the CaseViewer and CaseConverter softwares (3DHistech). For quantifications of immunofluorescence images, we used the Fiji ImageJ software (National Institute of Health) (Schindelin et al., 2012). In all EAE experiments, at least 2 sections and 2 areas per section were analyzed from each animal. In the case of cell cultures, at least 8 fields of view were analyzed per experiment. When z-stack projections were obtained from confocal microscopes, maximum projections were generated for the analysis. In the case of demyelinating lesions, lesion area was determined based on the absence of MBP staining and/or DAPI accumulation.

### Single nuclei RNA sequencing

For the study of the transcriptomic alterations caused by GABA_B1_ deletion in OPCs during EAE, single-nuclei RNA sequencing (snRNA-seq) was conducted. To this end, WT and cKO mice were perfused with PBS for 2 min, and spinal cords were collected and frozen at -80 °C until further processing. 1 WT and 1 cKO samples were included in the analysis, each of them consisting of a mixture of 1 male and 1 female spinal cord. For nuclei isolation, spinal cords were disrupted in homogenization buffer (10 mM Tris pH 8, 250 mM sucrose, 25 mM KCl, 5 mM MgCl2, 0,1 mM DTT, 40 U/μL RNasin Plus, 0.1 % triton X-100 v/v) using an automatic homogenizer. Next, tissue was incubated for 10 min on ice, with careful mixing every 2 min. The lysate was transferred to a glass homogenizer for further homogenization and then filtered through a 70 µm cell strainer, followed by two passess through 30 µm cell strainers. The resulting suspension was washed by centrifugation at 900 g and 4 °C for 10 min. Debris removal was performed through iodixanol gradient (iodixanol 29 %-iodixanol 50 %) centrifugation at 13,500 g and 4 °C for 20 min. Libraries were generated using the Chromium Next GEM Single Cell 3’ Kit v3.1 (10X Genomics #1000269), following the manufacturer’s instructions. Two samples (1 WT and 1 cKO) were sequenced on an Illumina Novaseq 6000 (National Genomics Infrastructure, Sweden) with a 28-10-10-91 read setup for RNA (minimum 20,000 read pairs per cell).

For the computational analysis, fastq files from both samples were processed using CellRanger v6.1.2. from 10X Genomics. Fastq files were input to the pipeline, and mm10 v2020-A-2.0.0 was used as reference genome. After alignment and demultiplexin, the resulting filtered feature-barcode matrix was imported into R software v4.2.1 for further processing.

For the subsequent computational analysis, we followed the workflow outlined by Seurat vignettes (Hao et al., 2021; Stuart et al., 2019). Initially, we evaluated the quality metrics for each sample, including the number of features/genes (nFeature), the number of counts (nCount) and the percentage of mitochondrial genes (percent-mito) per nucleus. We investigated both WT and cKO samples and established filtering thresholds accordingly: 50<nFeature<10,000, nCount<25,000 and percent-mito<5 for the WT and 50<nFeature<6,000, nCount<10,000 and percent-mito<5 for the cKO. The final dataset included 20,385 WT nuclei and 17,405 cKO nuclei for the downstream analysis.

Doublet detection was performed using DoubletFinder v2.0.3 (McGinnis et al., 2019), representing potential droplets with more than one nucleus. We marked any nuclei predicted as a doublet and, after a first inspection, we decided to remove them. Biological sex was predicted (female, F and male, M) using three features from the RNA assay for each nucleus, chrX-score, chrY-score and Xist-score. For the sex chromosome scores we retrieved the chrX and chrY from Ensembl annotations and calculated a global chrX and chrY-score. For the Xist-score, we used the AddModuleScore nbin = 2 function from Seurat. The prediction was then conducted using Classifier type Random forest from the Caret R package (Kuhn, 2008). The classifier was trained in a supervised manner in a control balanced subset of mouse spinal cord nuclei with known male and female classes, fitted model with 100 trees.

Then, we normalized the data applying log and Seurat canonical transform (SCT) normalization to the whole object (SCT data). We identified 2,000 highly variable features within the normalized dataset and then applied a linear transformation to scale the data. Next, we selected the first 40 principal components (PCs) of the data, as ranking through elbow plot proved that these included the main representation of the dataset. Sample integration was performed using Harmony on the selected PCs, using as grouping variables the WT and cKO samples. To better visualize our snRNA-seq data complexity, we applied the uniform manifold approximation and projection (UMAP) non-linear dimension reduction to these PCs, obtaining a 2D manifold. Unsupervised cell clustering of the whole dataset was conducted selecting a 0.1 resolution to obtain general cell types as the final clusters. The unsupervised clusters were annotated based on their cell type, according to their top 25 differentially expressed genes (Seurat FindAllMarkers function, default parameters).

In order to evaluate the differences in gene expression between WT and cKO, we subsetted OPCs and MOLs from the original dataset, and performed further downstream analysis on these two clusters. We followed the same workflow as for the original dataset, including normalization, data scaling, dimensional reduction and unsupervised cell clustering. Again, these unsupervised clusters were annotated based on their top 25 differentially expressed genes (Seurat FindAllMarkers function, default parameters). To obtain the differential expressed genes between WT and cKO, we carried out FindMarkers function of Seurat between both idents, setting an expression threshold of 1, and subsequently performed visualization and enrichment analysis with Seurat, ggplot2 (Wickham et al, 2016) and clusterProfiler (Wu et al, 2021; Yu et al, 2012). Processed data will be uploaded conviently at GEO repository (https://www.ncbi.nlm.nih.gov/geo/) after publication.

### Western Blot

For protein extraction from cultured oligodendrocytes, about 120,000 cells were first rinsed twice with cold phosphate-buffered saline (PBS) and then scrapped into RIPA buffer (Thermo Fisher #89900) supplemented with 0.05 M EDTA and protease inhibitor cocktail (Thermo Fisher #78440). Afterwards, cell lysates were diluted in sodium dodecyl sulfate (SDS) sample buffer (62.5 mM Tris pH 6.8, 10 % glycerol, 2 % SDS, 0.002 % bromophenol blue and 5.7 % β-mercaptoethanol, w/v) and stored at -20 °C. The entire process was carried out on ice. Samples were boiled at 100 °C for 8 min to ensure protein denaturation before loading into the electrophoresis gel.

Protein extracts from cell or tissue samples were separated by size through SDS-PAGE in 4-20 % (any kDa) Criterion TGX precast gels (Bio-Rad #5671124). Electrophoresis was conducted using a Tris-Glycine buffer (25 mM Tris, 192 mM glycine, 0.1 % SDS, pH 8.3), and the separated samples were then transferred to nitrocellulose membranes (Bio-Rad #1704159) using a Trans-Blot Turbo Transfer System (Bio-Rad). Membranes were blocked for 1 h at RT in blocking solution, consisting of Tris buffer saline (TBS, 20 mM Tris, 1.4 M NaCl, pH 7.6) with 0.05 % Tween-20 (v/v) (TBST), supplemented with 5 % bovine serum albumin (BSA, w/v) (Nzytech #MB04602). Afterwards, they were incubated with primary antibodies diluted in blocking buffer at 4 °C overnight. Primary antibodies were: mouse anti-GABAB1 (abcam #ab55051), mouse anti-MBP (1:1000, Biolegend #808402) and mouse anti-GAPDH (Millipore #MAB374). The next day, membranes were washed three times in TBST for 10 min at RT and were then incubated for 1 h at RT with horseradish peroxidase (HRP)-conjugated horse anti-mouse (Cell Signaling #7076) secondary antibody. Following three additional washes, protein bands were visualized using NZY Standard (Nzytech #MB40101) or NZY Advanced ECL (Nzytech #MB40201) to enhance the chemiluminescence signal in a ChemiDox XRS Imaging System (Bio-Rad). ImageLab software (Bio-Rad) was used for the quantification of protein band signal, and GAPDH was used for signal normalization.

### Flow cytometry

For the analysis of immune cell population during EAE development after baclofen treatment, cells from spleen and spinal cord were analyzed. Spleens and spinal cords were mashed through 70 µm strainers to create single cell suspensions. For the spinal cord, myelin debris was removed by a Percoll gradient centrifugation at 800 g for 10 min without break. In both cases, red blood cells were removed by incubation with lysis buffer (BD Biosciences, #349202) for 5 min at RT. Cell suspensions were washed with PBS containing 2.5 % BSA by centrifugation at 300 g for 5 min. Cells were then incubated with Fc blocker (1:500, BD Biosciences #553141) for 15 min at 4 °C and subsequently with fluorochrome-conjugated antibodies for 30 min at 4 °C. LIVE/DEAD^TM^ Fixable Near-IR Stain Kit (1:1000, Thermofisher #L10119) was included to identify and then exclude dead cells. The antibodies used for flow cytometry were: TCR γδ BV421 (1:100, BD Biosciences #562892), Ly6G BV510 (1:100, BD Biosciences #569685), CD4 BV605 (1:100, BD Biosciences #563151), CD3 BV650 (1:100, BD Biosciences #569683), CD45 APC (1:100, BD Biosciences #559864), CD11b (1:100, Biolegend #101205), CD8a (1:200, Biolegend #100707), Ly6C PercP Cy5.5 (1:100, Biolegend #128012), CD11c PE Cy7 (1:100, Biolegend #117318). After three washes in PBS, samples were acquired in a LSR Fortessa X-20 flow cytometer (BD Biosciences) and analyzed with FlowJo v.10.8.1 (Tree Star) software.

### Statistical analysis

Data are presented as mean ± standard deviation (SD), and dots represent the number of animals analyzed or independent experiments, as specified in each figure legend. All efforts were made to maintain blinded conditions throughout the experiments. Sample size was determined according to previous experimental approaches established in the laboratory. Statistical analyses were conducted using GraphPad Prism 8 software, applying the appropriate test for each experiment, as outlined in the figure legends. Normality of the data was assessed using Anderson-Darling, D’Agostino and Pearson, Shapiro-Wilk and Kolmogorov-Smirnov tests. In general, for comparisons between two groups, the analysis was performed using Student’s two-tailed t-test. In all cases, p-values < 0.05 were considered as statistically significant.

## Acknowledgements

This work was supported by grants from the Spanish Ministry of Science and Innovation (PID2022-140236OB-100 M.V.S-G; PID2022-143020OB-100 C.M); BioEF Foundation (BIO23/EM/007 M.V.S-G), the Basque Government (IT1551-22, PIBA-2024-1-0037, and 2024333031 M.V.S-G) and CIBERNED (CB06/05/00 76 C.M). L.B-C hold a predoctoral fellowship from the Basque Government; B.I.O-B from the Spanish Ministry of Education and Science; I.L-E from the University of the Basque Country UPV/EHU and R.S a neuroscience predoctoral fellowship from Fundación Tatiana. L.A is an Ikerbasque Research Fellow (COFUND program H2020-MSCA-COFUND-2020 101034228-WOLFRAM2) and F.B is an Ikerbasque Research Professor. The computation analysis and data handling was enabled in part by resources provided by the National Academic Infrastructure for Supercomputing in Sweden (NAISS), partially funded by the Swedish Research Council through grant agreement no. 2022-06725. The authors thanks to the SGIker platform from the University of the Basque Country (UPV/EHU) and the facilities at Achucarro Basque Center for Neuroscience for the technical assistance.

---

**Extended figure 1.**
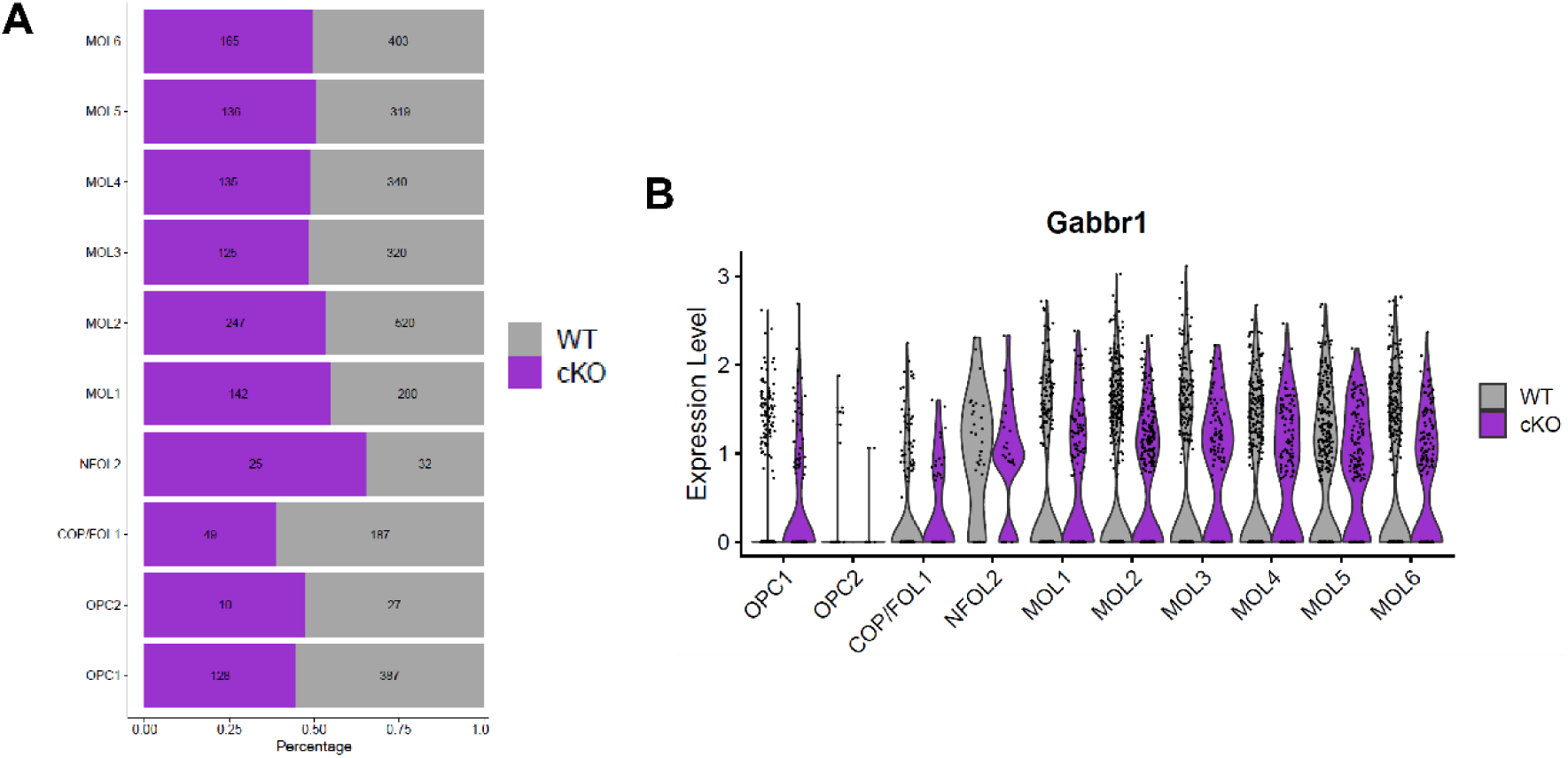
*Gabbr1* expression along oligodendrocyte subtypes in NG2-GABA_B1_ cKO. (A) Histogram showing the percentage of nuclei distributed among oligodendrocyte subtypes in WT (grey) and cKO (purple) mice. (B) Violin plot showing the relative expression of *Gabbr1* gene across oligodendrocyte subtypes.

**Extended figure 2.**
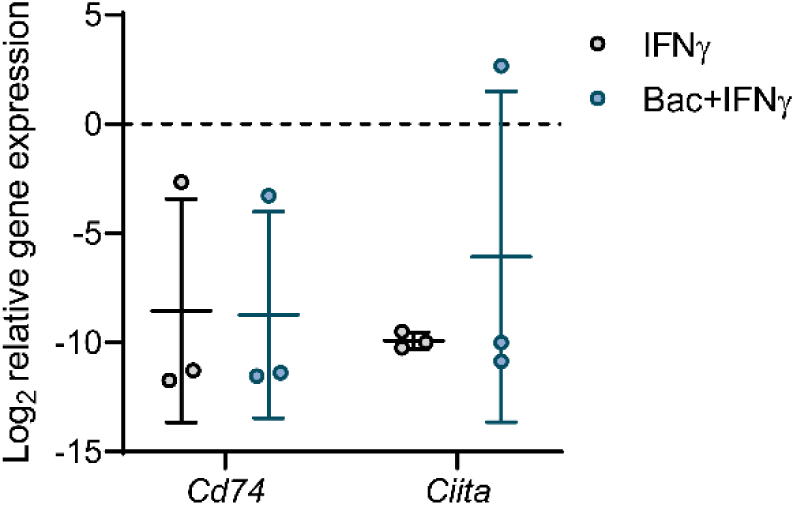
Baclofen treatment does not modify oligodendrocyte immune gene expression induced by IFNγ. Histogram showing fold changes in the expression of *Cd74and Ciita* genes relative to control in cortical OPCs treated for 3 DIV with baclofen. Values are determined as log_2_ and data were corrected using the geometric mean of the reference genes *Ppia* and *Pgk1*. Relative expression values were calculated with the 2^-ΔΔCt^ method and are expressed as relative to control (untreated cells; value 0). Paired one-way ANOVA. Data represent mean ± SD and dots represent independent experiments.

**Extended figure 3.**
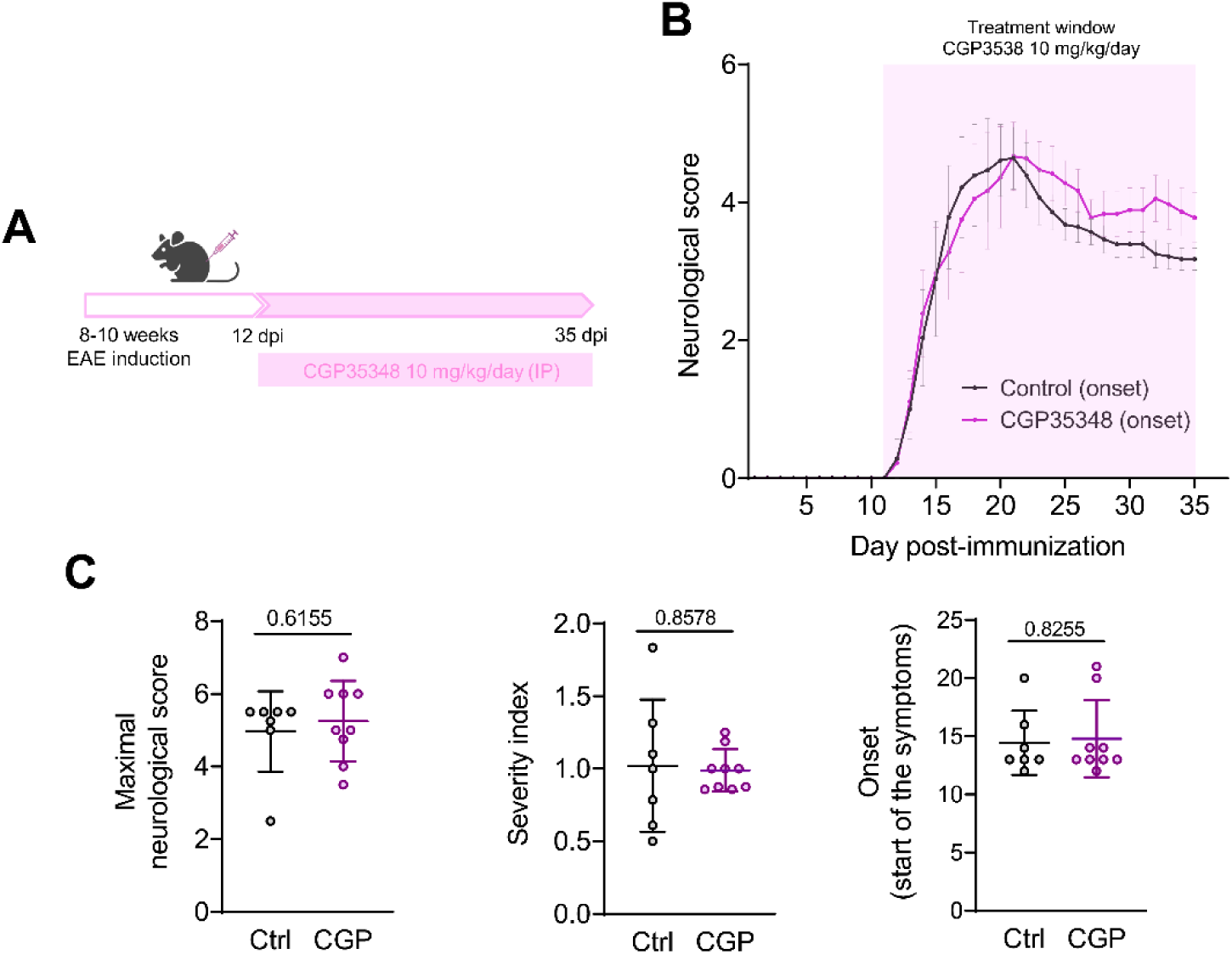
CGP35348 treatment does not significantly alter disease course in the EAE model. (A) Diagram showing the experimental setup for CGP35348 (CGP) administration in the EAE model. (B) Progression of the neurological score relative to motor symptoms in control and CGP-treated mice, with the treatment starting from the onset of the symptoms. Data represent mean ± SD from at least 7 animals. (C) Histograms showing the maximal score, severity index and onset of the symptoms for every animal. Unpaired Student’s t-test. Data represent mean ± SD and dots represent different animals.

